# Associations between regional blood-brain barrier disruption, aging, and Alzheimer’s disease biomarkers in cognitively normal older adults

**DOI:** 10.1101/2024.02.16.580788

**Authors:** Marisa Denkinger, Suzanne Baker, Ben Inglis, Sarah Kobayashi, Alexis Juarez, Suzanne Mason, William Jagust

## Abstract

**Background:** Blood-brain barrier disruption (BBBd) has been hypothesized as a feature of aging that may lead to the development of Alzheimer’s disease (AD). We sought to identify the brain regions most vulnerable to BBBd during aging and examine their regional relationship with neuroimaging biomarkers of AD.

**Methods:** We studied 31 cognitively normal older adults (OA) and 10 young adults (YA) from the Berkeley Aging Cohort Study (BACS). Both OA and YA received dynamic contrast-enhanced MRI (DCE-MRI) to quantify K_trans_ values, as a measure of BBBd, in 37 brain regions across the cortex. The OA also received Pittsburgh compound B (PiB)-PET to create distribution volume ratios (DVR) images and flortaucipir (FTP)-PET to create partial volume corrected standardized uptake volume ratios (SUVR) images. Repeated measures ANOVA assessed the brain regions where OA showed greater BBBd than YA. In OA, K_trans_ values were compared based on sex, Aβ positivity status, and APO*E4* carrier status within a composite region across the areas susceptible to aging. We used linear models and sparse canonical correlation analysis (SCCA) to examine the relationship between K_trans_ and AD biomarkers.

**Results:** OA showed greater BBBd than YA predominately in the temporal lobe, with some involvement of parietal, occipital and frontal lobes. Within an averaged ROI of affected regions, there was no difference in K_trans_ values based on sex or Aβ positivity, but OA who were APO*E4* carriers had significantly higher K_trans_ values. There was no direct relationship between averaged K_trans_ and global Aβ pathology, but there was a trend for an Aβ status by tau interaction on K_trans_ in this region. SCCA showed increased K_trans_ was associated with increased PiB DVR, mainly in temporal and parietal brain regions. There was not a significant relationship between K_trans_ and FTP SUVR.

**Discussion:** Our findings indicate that the BBB shows regional vulnerability during normal aging that overlaps considerably with the pattern of AD pathology. Greater BBBd in brain regions affected in aging is related to APOE genotype and may also be related to the pathological accumulation of Aβ.

## Introduction

Brain aging is accompanied by the aggregation of pathological proteins and the increasing prevalence of cerebrovascular disease. Recent research has shown that blood brain barrier disruption (BBBd) is an important feature of both brain aging and Alzheimer’s disease (AD). BBBd in human aging and AD has been documented through the detection of blood-derived proteins in the hippocampus (HC) and cortex of AD patients and increases in the cerebrospinal fluid (CSF) of the plasma albumin protein ratio (Qalb) in both aging and AD [1–4]. More recent evidence of BBBd in humans comes from studies using the high spatial and temporal resolution imaging technique, dynamic contrast-enhanced magnetic resonance imaging (DCE-MRI), which allows measurement of subtle BBB changes [5]. A number of studies using DCE MRI have shown BBBd in both aging and AD with particular vulnerability of the hippocampus to this process [6–12]. Major questions remain, however, regarding the overall spatial distribution of BBBd, whether abnormalities are limited to the medial temporal lobe (MTL) and most importantly, whether or how BBBd is related to the development of AD.

Studies that explore the relationship between BBBd and AD benefit from the availability of fluid and PET biomarkers of the two protein aggregates associated with the disease –β-amyloid (Aβ) and pathological forms of tau. BBBd measured with DCE-MRI in the HC and parahippocampal cortex (PHC) is evident before CSF measures of AD pathology, or cognitive decline [8]. Some evidence also suggests a lack of association between BBBd, measured using Qalb, and global Aβ PET in non demented older adults [1]. A recent study using DCE-MRI in cognitively normal and impaired individuals reported a lack of association of BBBd with Aβ or tau positivity, but a relationship with cognitive impairment and APO*E4* genotype [13]. The discordance between in vivo measurement of BBBd and evidence of AD pathology suggests that BBBd may either be a very early precursor of AD, or lead to dementia symptoms through a mechanism independent of amyloid and tau pathology [14].

In this study, we investigated the relationship between BBBd and AD through 2 lines of evidence. First, we examined the full spatial distribution of BBBd which offers an ability to draw inferences about causal mechanisms and to help establish the role of BBBd in dementia. To do this, we compared BBB function in a group of cognitively normal older adults (OA) to young adults (YA) and mapped the whole brain distribution of BBBd. Second, we investigated whether BBBd in OA was associated with APO*E4* genotype and regional Aβ and tau, measured using PET imaging.

## Methods

### Participants

We recruited 31 cognitively normal OA and 10 YA enrolled through the Berkeley Aging Cohort Study (BACS). OA participants were part of an ongoing longitudinal study of aging and received neuropsychological testing, DCE-MRI, and both Aβ and tau PET. We acquired PET scans an average of 2.4 months (SD=5.3) before or after the DCE-MRI. YA participants received neuropsychological testing and DCE-MRI only. Inclusion criteria included a baseline Mini Mental State Examination (MMSE) score of >26, scores on all neuropsychological tests within 1.5 SD of age, sex and education adjusted norms, no neurological, psychiatric, or major medical illness, and no medications affecting cognitive ability.

### Standard Protocol Approvals, Registrations, and Participant Consents

The project was approved by the institutional review board (IRB) at the University of California, Berkeley, and written informed consent was collected from each participant. The recruitment period for this study was 12/16/2019 – 6/23/2023.

### MRI Acquisition

1.5T MRI data were collected for standard PET processing at the Lawrence Berkeley National Laboratory (LBNL) on a Siemens Magnetom Avanto scanner. A whole-brain high resolution sagittal T1-weighted MPRAGE scan was acquired for each participant (TR= 2110 ms, TE=3.58 ms, voxel size= 1mm isotropic, flip angle= 15°).

3T MRI data were collected at the UC Berkeley Henry H. Wheeler, Jr. Brain Imaging Center with a 3T Siemens Trio scanner and 32-channel head coil. High resolution sagittal T1-weighted magnetization prepared rapid gradient echo (MPRAGE) scans were acquired for each participant (repetition time (TR)=2300 ms, inversion time (TI)=900 ms, echo time (TE)=2.96 ms, flip angle=9°, voxel size=1 mm isotropic, field of view (FOV)=256 x 240 x 176 mm).

Baseline coronal T1-weighted maps were acquired using a T1-weighted three-dimensional (3D) spoiled gradient echo pulse sequence and variable flip angle method (TR=3.8 ms, TE=1.64 ms, voxel size=2.5 x 1.3 x 2.5 mm, FOV=240 x 180 x 220 mm, flip angles=2, 5,10,12,15°) with full brain coverage. Coronal DCE-MRI were acquired with the same sequence and a flip angle of 5°. The sequence was repeated for a total of 21.3 minutes with a time resolution of 18 seconds [15]. The macrocyclic gadolinium-based contrast agent Gadobutrol (gadavist, 1 mmol/ml, 0.1ml/kg body weight) was administered intravenously over 30 seconds following the first 7 DCE repetitions.

### PET Acquisition

Methods for PET acquisition and analysis have been described previously, but are summarized here [16]. All PET scans were acquired at LBNL on a Siemens Biograph PET/CT scanner with the radiotracers [^11^C]PiB for Aβ and [^18^F]Flortaucipir (FTP) for tau synthesized at LBNL’s Biomedical Isotope Facility. Following acquisition of a CT scan, PiB-PET data were collected across 35 dynamic acquisition frames for 90 minutes after injection and FTP-PET data were binned into 4 x 5 minute frames from 80-100 minutes after injection. All PET images were reconstructed using an ordered subset expectation maximization algorithm, with attenuation correction, scatter correction, and smoothing with a 4 mm Gaussian kernel.

### Structural MRI Processing

The 3T T1-weighted images were segmented using FreeSurfer v7.1.1 (http://surfer.nmr.mgh.harvard.edu/) to derive anatomical ROIs in native space for the measurement of BBBd. Segmentations and parcellations were visually checked to ensure accuracy. FS Desikan-Killiany atlas ROIs were extracted and used to calculate region-specific BBBd. The 1.5T MRIs were used only for PET coregistration and were segmented in our standard processing pipeline to derive native space FS ROIs for PiB and FTP quantification.

### DCE-MRI Processing

DCE-MRI scans were realigned to the first image for motion correction using Statistical Parametric Mapping 12 (SPM12) prior to analysis. Analysis was completed using the DCE-MRI analysis software, ROCKETSHIP, running with Matlab [17]. The arterial input function (AIF) was manually labeled in each participant at the common carotid artery and was fitted with a bi-exponential function prior to kinetic modeling. A modified version of the Patlak linearized regression mathematical analysis was used to generate BBB permeability volume transfer constant (K_trans_) maps [18]. This model provides high accuracy and precision for small permeability values [19, 20]. The total contrast agent concentration in the brain tissue, C_tissue_ (*t*), can be described as a function of the contrast agent concentration in plasma, CAIF (*t)*, the volume fraction of plasma, *v*p, and the blood-to-brain volume transfer constant, K_trans_, using the following equation:

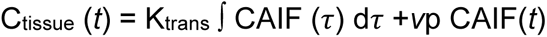

K_trans_ thus represents the transport from the intravascular space to the extravascular extracellular space, with a higher K_trans_ indicating greater BBBd. K_trans_ was calculated at a voxelwise level for each subject and was averaged within FS Desikan-Killiany atlas ROIs.

### PET Processing

FTP images were realigned, averaged, and coregistered to the participant’s 1.5T MRI using SPM12. Standardized uptake value ratio (SUVR) images were calculated by using the mean tracer uptake from 80-100 minutes post-injection and were normalized with an inferior cerebellar gray reference region [21]. The average SUVR values were calculated in FS Desikan-Killiany atlas ROIs derived from segmentation of the participant’s MRI. This ROI data was partial volume corrected (PVC) using a modified Geometric Transfer Matrix approach and these values were used for analyses [21, 22].

PiB images were also realigned using SPM12. The frames from the first 20 minutes of acquisition were used for coregistration to the participant’s 1.5T MRI. Distribution volume ratio (DVR) images for the PiB frames 35-90 minutes post injection were calculated using Logan Graphical analysis [23] and using whole cerebellar gray as a reference region. Global Aβ uptake was calculated using FS cortical ROIs [24]. A DVR of greater than1.065 was used to classify participants as Aβ+ [25]. We also calculated the mean DVR within a set of FS Desikan-Killiany atlas ROIs that reflect the typical pattern of Aβ deposition.

### Statistical Analyses

K_trans_ values were not normally distributed, as indicated by the Shapiro-Wilk’s test and were therefore log transformed. Statistical analyses were conducted using jamovi (https://www.jamovi.org/) and RStudio version 4.2.3 (https://www.rstudio.com/). To investigate regional differences in K_trans_ values between OA and YA, we ran a repeated measures analysis of variance (ANOVA) (group, region, and group X region), followed by post-hoc independent sample *t*-tests. S1 Table lists the 37 ROIs used in this analysis. We ran the analysis using K_trans_ values averaged across hemispheres, followed by right and left hemispheres comparisons in the OA. Subsequent analyses examining relationships between K_trans_ and AD biomarkers used only those regions where OA showed significantly larger K_trans_ values than YA. First, we created an averaged K_trans_ variable using these 20 regions and reported comparisons between sex, Aβ status, and APO*E4* carrier status. We also ran a linear model predicting averaged K_trans_ from EC FTP and an EC FTP by Aβ status interaction, controlling for age and sex. We also ran the same model using FTP in a temporal meta-ROI [26].

Sparse Canonical Correlation Analysis (SCCA) was used to examine the multivariate regional relationships between BBBd and AD biomarkers, including PiB and FTP. SCCA is a variant of the traditional Canonical Correlation Analysis (CCA), which finds the optimal linear combinations of variables from two different modalities that are highly correlated with each other by weighting each variable to determine its significance in the correlation [27, 28]. The original variables are multiplied by these weights to form a multivariate projection. The canonical correlation is the correlation between these multivariate projections and multiple canonical correlations can be derived by using the residual data of the canonical variates to compute the subsequent canonical correlation. The first canonical correlation is usually the highest, capturing the maximum possible correlation between the two sets of variables. SCCA enhances CCA by incorporating sparsity constraints into the canonical vectors, often through penalties like the Lasso penalty. This process ensures that many weights in the canonical vectors are zero, highlighting the most significant variables in each modality contributing to the correlation.

We performed SCCA using the using the Penalized Multivariate Analysis (PMA) R package [29]. Bilateral K_trans_, FTP and PiB ROIs were chosen for analysis to reduce redundancy in the model. We used the 20 ROIs that were most affected by aging in our sample. The effects of age and sex were removed from the data by calculating the residuals, which were used for analyses. A lasso penalty of 0.5 was used to extract the most meaningful ROIs and we did not constrain any of the weights in the model to be positive or negative. The data were also mean centered and scaled. SCCA was performed separately for K_trans_ and PiB and K_trans_ and FTP. Significance was determined by correlating the multivariate projections of K_trans_ and AD biomarkers (PiB, FTP), which produces a correlation coefficient in each dimension. An F-approximation of Wilk’s lambda was used as a test statistic and p-values < 0.05 were considered significant.

### Data Availability

Data will be made publicly available to qualified investigators following publication of this study.

## Results

### Participant Characteristics

The sample consisted of 31 OA (68-85 years old, mean 77.5, SD 5.2) and 10 YA (22-28 years old, mean 24.3, SD 2.3). Table 1 shows the sample characteristics for OA and YA. Groups did not differ on years of education or sex. Within the OA group, 14 participants were classified as Aβ+, 4 as *APOE4* carriers, and 8 had treated hypertension (HTN). The range of partial volume corrected FTP values in the entorhinal cortex was 0.8-1.9 and in the temporal meta-ROI was 1.0-2.4.

**Table 1.** Participant Demographics. Abbreviations: MMSE = Mini-Mental State Examination; APO*E4* = Apolipoprotein ε4; HTN = Hypertension. DVR = Distribution volume ratio; EC = Entorhinal cortex; PVC = Partial volume corrected; SUVR = Standardized uptake value ratio Data shown as mean ± SD for continuous variables or n (%) for categorical variables. Group comparisons were run using independent sample *t-*tests or *χ2* tests. * *p* < 0.05 ** *p* < 0.001

|  | Old (n = 31) | Young (n = 10) |
| --- | --- | --- |
| **Age, y** ** | 77.5 ± 5.2 | 24.3 ± 2.3 |
| **Female** | 16 (52%) | 3 (30%) |
| **Years of education** | 17.6 ± 1.7 | 18.6 ± 2.1 |
| **MMSE** * | 28.4 ± 1.4 | 29.6 ± 0.9 |
| **APOE4 carriers** | 4 (14%); n = 28 | n/a |
| **Aβ+** | 14 (45%) | n/a |
| **Global PiB (DVR)** | 1.1 ± 0.2 | n/a |
| **EC FTP (PVC SUVR)** | 1.4 ± 0.3 | n/a |
| **Temporal meta-ROI FTP (PVC SUVR)** | 1.4 ± 0.2 | n/a |

### Generation of K_trans_ Maps

Fig 1 shows example whole brain K_trans_ maps in 2 OA and 2 YA participants with high and low BBBd (defined as the highest and lowest average K_trans_ from each group in averaged temporal, parietal, and occipital lobes). The 2 OA participants demonstrated larger K_trans_ values distributed throughout the cortex than the YA, compared to limited and more localized BBBd.

**Fig 1.**
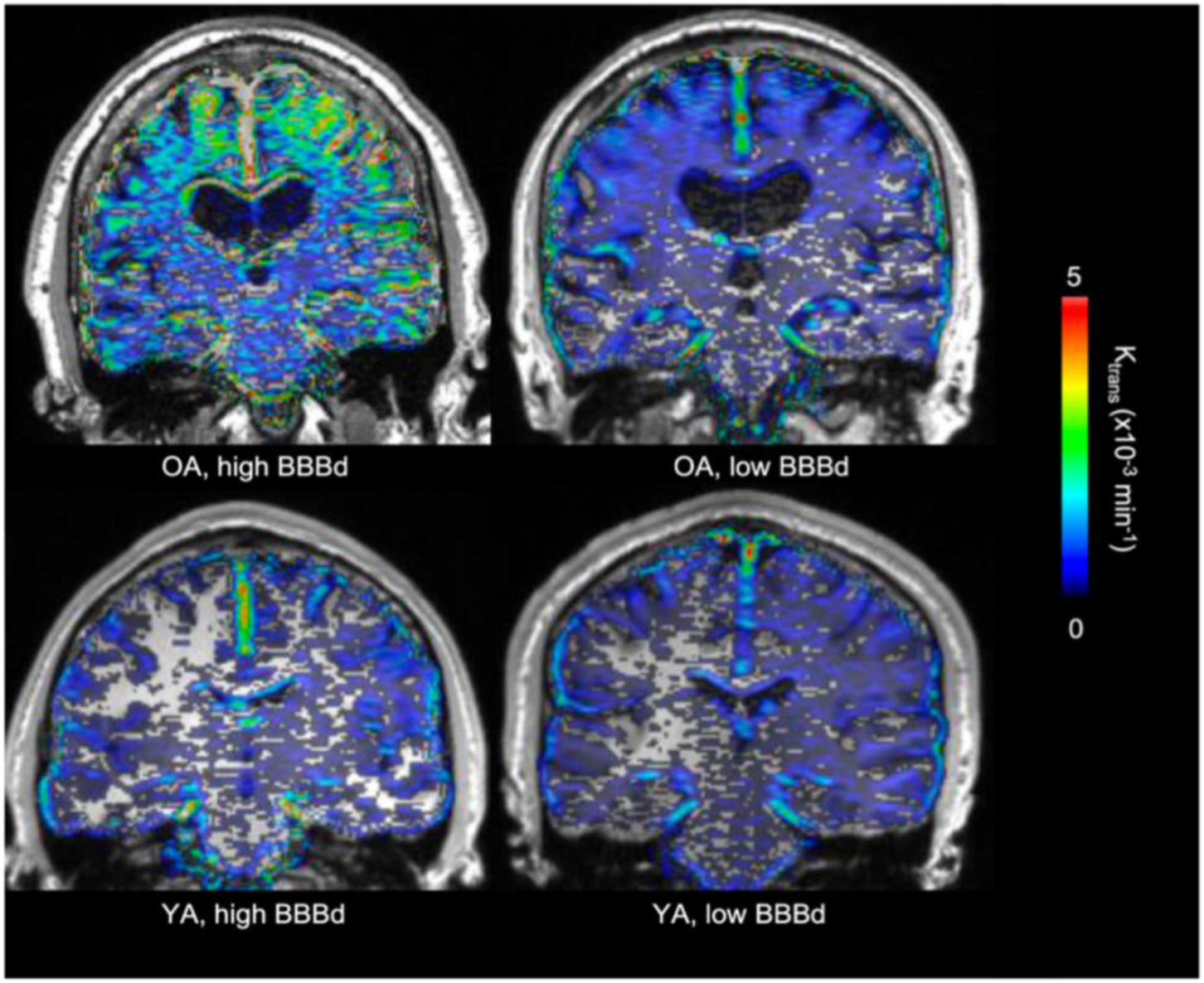
Voxelwise K_trans_ Maps. Representative voxelwise BBB K_trans_ maps in 2 OA (top) and 2 YA (bottom) participants classified as having high and low BBBd. These are the raw K_trans_ values with units of min^-1^.

### Differences in BBBd Between OA and YA in Cortical Regions

A repeated measures ANOVA (age group, region, age group X region) across the BBBd average calculated in 37 bilateral FS-defined cortical ROIs showed a significant main effect of region and an age group by region interaction (F = 2.1, p<0.001). Post-hoc independent sample *t*-tests showed that OA had larger K_trans_ than YA in 20 regions bilaterally, that were largely in temporal and parietal cortex: (amygdala (Amyg), banks of the superior temporal sulcus (BanksSTS), entorhinal cortex (EC), fusiform gyrus (Fu), hippocampus (HC), insula (Ins), inferior temporal (IT), middle temporal (MT), parahippocampus (PHC), transverse temporal (TrT)), parietal (inferior parietal (IP), isthmus of the cingulate gyrus (IstCg), posterior cingulate (PCC), precuneus (PreCu)), and occipital lobes (cuneus (Cu), lateral occipital (LO), lingual (Lg), pericalcarine (PerCa)). There were also two regions in the frontal lobe where OA had larger K_trans_ (pars opercualris (Op), paracentral (PaC)) (S1 Fig, all *p* < 0.05). Only the PHC and IstCg survived corrections for multiple comparisons (p < 0.001). We did not find any other regions with significant K_trans_ differences between groups, including white matter, nor were there any ROIs where YA had higher K_trans_ than OA. Effect sizes for the left and right hemisphere separately are shown in Fig 2. We used a paired samples *t*-test to compare left and right hemisphere K_trans_ values in the OA and found significantly larger K_trans_ in the left bankssts, Fu, HC, Ins, IP, IstCg, Lg, MT, PerCa, PHC, and TrT, with the bankssts, Fu, IstCg, IP, and PHC surviving corrections for multiple comparisons (*p* < 0.001). The overall pattern of increased BBBd was similar across hemispheres, so we used averaged bilateral data for the rest of the analyses to reduce the number of ROIs.

**Fig 2.**
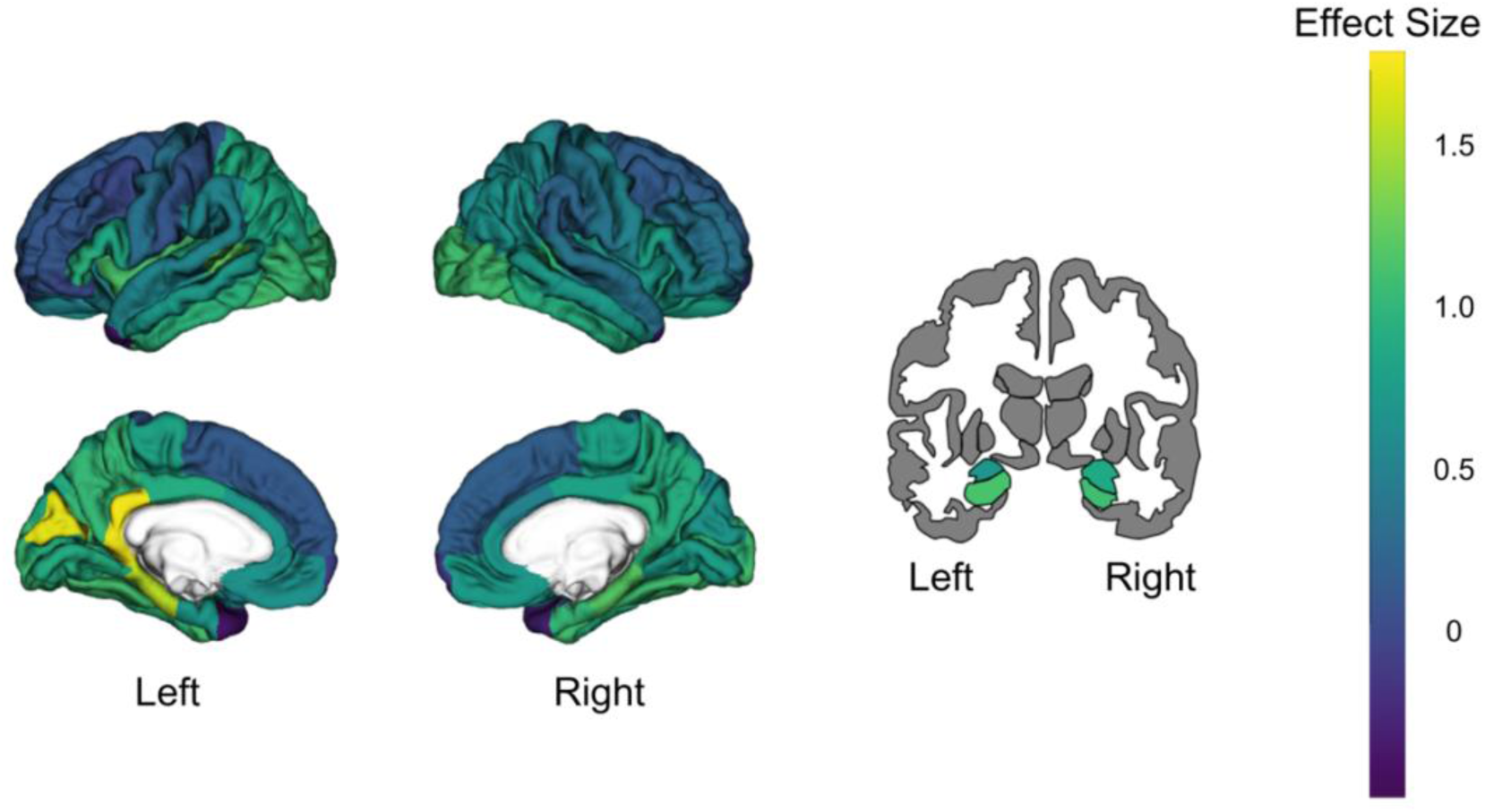
K_trans_ Differences Between OA and YA. Results from independent sample *t-*test comparing log transformed K_trans_ values in 74 FS ROIs between OA and YA. Brain plots show the Cohen’s d effect size for each ROI. Effect sizes were overall larger in the left hemisphere than the right hemisphere.

### Averaged BBBd Comparisons and Correlations

We created an averaged K_trans_ ROI consisting of the 20 regions where OA showed significantly larger K_trans_ than YA. Within the OA participants, there was no significant correlation between age and the averaged K_trans_ ROI (*R* = 0.22, *p* = 0.34). There were no significant averaged K_trans_ differences by sex (*t =* −0.75, *p* = 0.46) or Aβ status (*t* = −1.27, *p* = 0.21). There was also no significant relationship between averaged K_trans_ and global PiB index (*R =* 0.32, *p* = 0.22). We did find a significant difference by APO*E4* status (*t = −2.50, p* = 0.02), where APO*E4* carriers had greater K_trans_. There was no significant main effect of EC FTP or temporal meta-ROI FTP on predicting averaged K_trans_, but there was trend level interaction for EC FTP and Aβ status (*R* = 0.41, *p* = 0.08), as well as meta-ROI FTP and Aβ status (*R =* 0.42, *p* = 0.08).

### Regional relationships between BBBd and AD biomarkers

Figs 3 and 4 show visual representations of the relationships between K_trans_ and AD biomarkers for the first three canonical correlation dimensions, and S2 and S3 Table show the weights of each region that contributed to the dimension.

**Fig 3.**
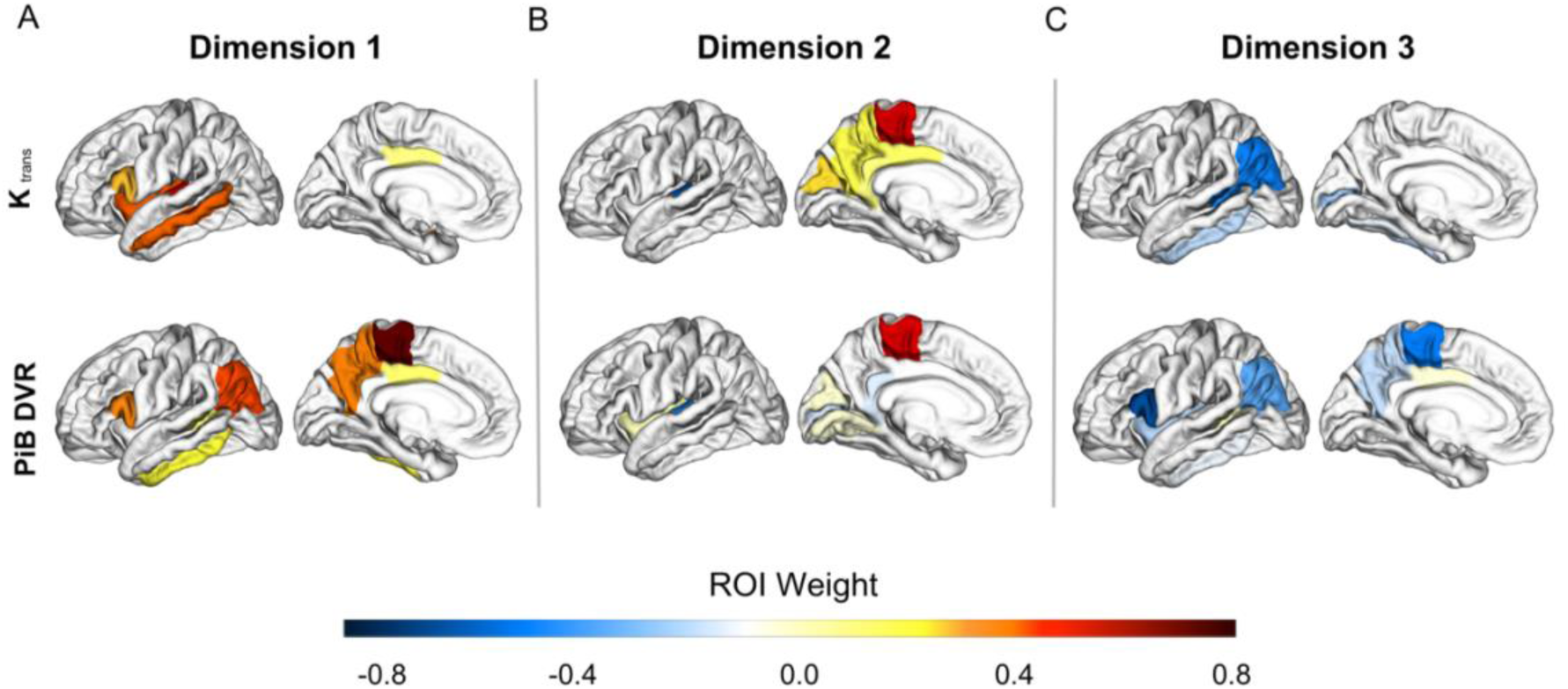
Associations Between BBBd and Aβ. Brain plots show the first three dimensions from the sparse canonical correlation analysis between K_trans_ and PiB ROIs controlled for the effects of age and sex (A-C). (A) Dimension 1, *r* = 0.39, F (9, 61) = 2.8, *p* = 0.009 (B) Dimension 2, *r* = 0.67, F (4, 52) = 5.2, *p* = 0.001 (C) Dimension 3, *r* = 0.26, F (1, 27) = 2.0, *p* = 0.17. Dimensions represent K_trans_ changes (increase or decrease) aligned with corresponding PiB changes. Increases in a variable are signified by positive weights and decreases with negative weights. Regions are colored based on their weight and bilateral ROIs are depicted on a left hemisphere template brain. Weights reduced to zero due to sparsity constraints are not included in the color scale.

**Fig 4.**
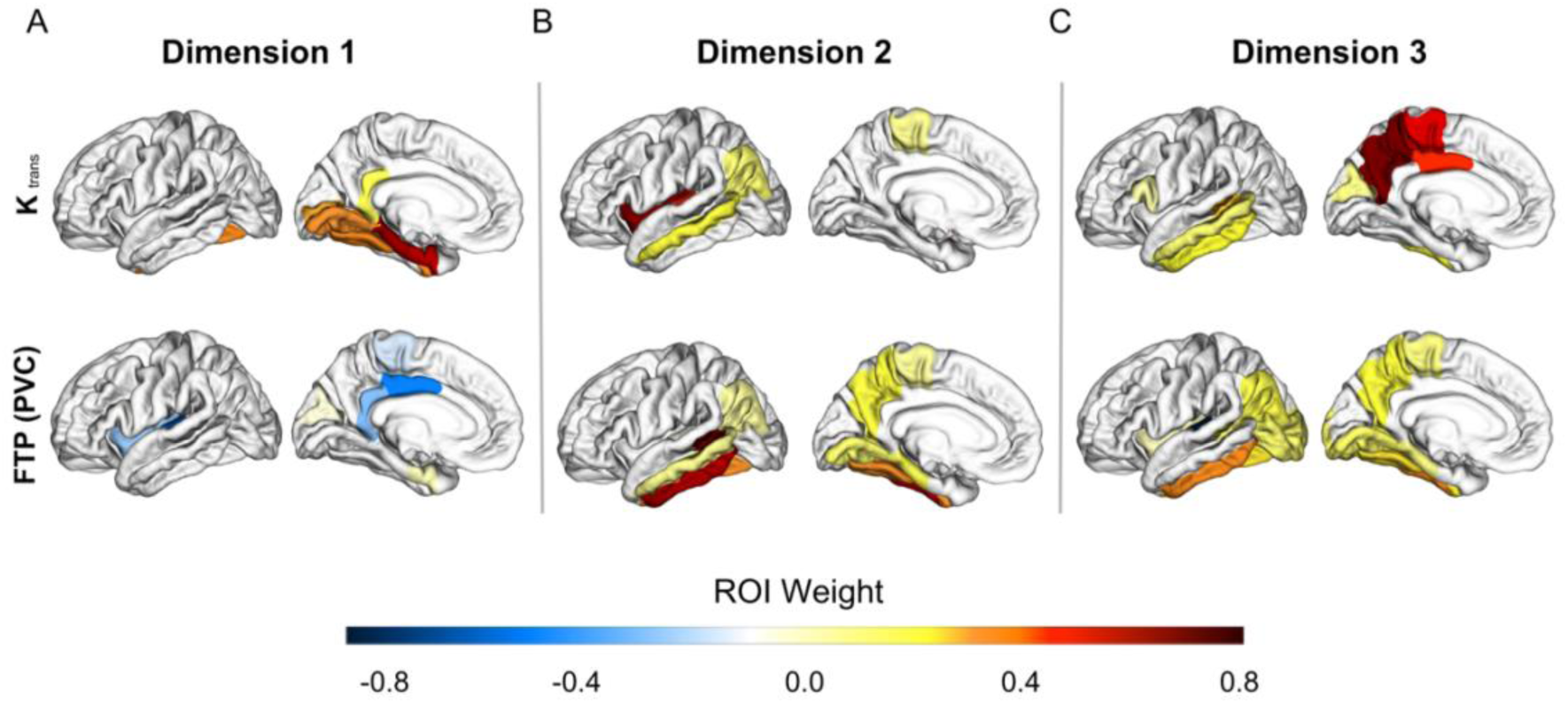
Associations Between BBBd and Tau. Brain plots show the first three dimensions from the sparse canonical correlation analysis between K_trans_ and partial volume corrected FTP ROIs controlled for the effects of age and sex (A-C). (A) Dimension 1, *r =* 0.42, F (9, 61) = 1.5, *p* = 0.16 (B) Dimension 2, *r* = 0.30, F (4, 52) = 2.1, *p* = 0.09 (C) Dimension 3, *r* = 0.43, F (1, 27) = 6.3, *p* = 0.02. Dimensions represent K_trans_ changes (increase or decrease) aligned with corresponding FTP changes. Increases in a variable are signified by positive weights and decreases with negative weights. Regions are colored based on their weight and bilateral ROIs are depicted on a left hemisphere template brain. Weights reduced to zero due to sparsity constraints are not included in the color scale.

The first two canonical correlations between K_trans_ and PiB were significant. Dimension 1 was represented by positive K_trans_ weights primarily in the temporal lobe and positive PiB weights in temporal and parietal cortices (Fig 3A, *r* = 0.39, F (9, 61) = 2.8, *p* = 0.009), indicating brain regions where higher PIB DVR was associated with higher K_trans_. Dimension 2 was represented by both positive and negative K_trans_ weights in parietal, occipital, and temporal cortices and mainly negative PiB weights in the temporal lobe (Fig 3B, *r* = 0.67, F (4, 52) = 5.2, *p* = 0.001). The dimension 3 correlation was not statistically significant (Fig 3C, *r* = 0.26, F (1, 27) = 2.0, *p* = 0.17).

Next, we looked at the associations between K_trans_ and partial volume corrected FTP in 20 regions. The correlation between K_trans_ and FTP in dimension 1 was not statistically significant (Fig 4A, *r =* 0.42, F (9, 61) = 1.5, *p* = 0.16). Dimension 2 showed a trend level correlation represented by positive K_trans_ weights mainly in the temporal lobe and positive FTP weights in the temporal and parietal cortices (Fig 4B, *r* = 0.30, F (4, 52) = 2.1, *p* = 0.09). Dimension 3 was represented by positive K_trans_ weights in lateral temporal and medial parietal regions and positive FTP weights in similar regions (Fig 4C, *r* = 0.43, F (1, 27) = 6.3, *p* = 0.02).

## Discussion

Better characterization of BBBd during aging and its relationship, if any, to AD biomarkers is critical in understanding the role of neurovascular dysfunction in the AD pathological cascade. Using DCE-MRI in cognitively normal OA and YA, we showed that BBBd does not occur globally, but rather occurred predominately in the temporal lobe, with involvement of the parietal, and less involvement of occipital and frontal lobes. In these regions we also found that APO*E4* carriers had greater BBBd than non-carriers. PET imaging showed that BBBd has weak and inconsistent relationships with AD pathology. Although the large group of brain regions with elevated BBBd did not show any relationship to Aβ, there was a trend for an Aβ by tau interaction on K_trans_ in this region, and the SCCA showed a pattern of regional relationships between K_trans_ and PiB DVR that recapitulated the known topography of AD pathology. Overall, these findings indicate that BBBd during aging occurs in overlapping regions affected in AD, is related to APOE genotype, and that it may be related to Aβ pathololgy.

The regional BBBd we found strikingly reflects the pattern of brain vulnerability to AD pathology, particularly in regions that are affected early. Tau accumulation in normal aging begins in the medial temporal lobe and spreads to neighboring regions in the inferolateral temporal and medial parietal lobes in the presence of Aβ [16, 30]. The pattern of brain Aβ accumulation overlaps with the spatial location of tau best in later disease stages, covering regions in prefrontal, parietal, lateral temporal, and cingulate cortices. In line with previous studies [6–9, 12, 13], we saw greater BBBd in the MTL, particularly the EC, PHC, and HC, which accumulate tau pathology and undergo atrophy in normal aging, but do not typically accumulate Aβ at early stages of AD [31]. We also saw that in our sample the frontal lobe is relatively spared from BBBd, which is interesting because this brain region is associated with early Aβ accumulation [31], but late tau accumulation [32]. These differences suggest that BBBd follows a distribution pattern more like tau accumulation than Aβ, with involvement of the MTL, temporal, parietal, and occipital lobes.

We next aimed to untangle the relationships between BBBd and neuroimaging measures of AD biomarkers. We found no significant difference in BBBd based on Aβ status in the prespecified ROIs, although there was a significant APO*E4* effect. We did not find any main effects of global Aβ, or regional tau in EC or temporal meta-ROI in predicting averaged K_trans_. However, we did find trend level interactions between Aβ status and tau, which suggests the possibility that the combined pathologies, which reflect the presence of AD, are related to BBBd. To further investigate the regional relationship between BBBd and AD biomarkers, we used a data driven SCCA approach. This method has the advantage of not requiring prespecified ROIs, and therefore may be able to detect subtle regional relationships. We observed that increased BBBd was associated with increased Aβ in temporal and parietal cortex, brain regions typically affected by Aβ pathology. However, the spatial relationships between tau pathology and BBBd revealed through this statistical approach were weak. Altogether, we interpret our results as pointing towards complex relationships between Aβ, tau and BBBd such that Aβ and BBBd could promote tau deposition over time, or Aβ and tau together could promote BBBd. Larger samples and longitudinal data will be necessary to establish these relationships.

The current evidence for a relationship between AD pathology and BBBd is conflicting. Existing studies use different methods for defining BBBd, and different ways of measuring AD pathology. Our findings of an effect of APO*E4* genotype on BBBd replicate results of one study; this study did not find any consistent or trend level relationships between DCE-measured BBBd and Aβ or tau pathology measured with PET, but did find an APOE effect [13]. A previous study using MRI measures of water exchange to characterize BBBd showed an association between greater BBB permeability in frontal, parietal, and temporal regions, and evidence of Aβ accumulation, measured as reduced CSF Aβ42 levels [33]. In a sample of patients with dementia, greater BBBd, measured using Q_alb_, was associated with less CSF Aβ42 and Aβ40, but was not associated with CSF pTau181 or tTau [34]. However, another study using MRI measures of water exchange, found that greater BBB permeability was associated with increased CSF pTau [35]. These studies are difficult to compare to one another because of the methodological differences but suggest the possibility of relationships between AD pathology and alterations in BBB function.

Associations between BBBd and AD pathology have also been probed with animal models, which also can assess temporal relationships. Previous research found that BBBd leads to the deposition of Aβ by increasing its production and preventing its normal transport across the BBB [36, 37]. Studies in animal models have also shown that BBB permeability is increased before the presence of Aβ pathology in an AD mouse model [38] and that loss of pericytes increased brain Aβ40 and Aβ42 levels [39]. In contrast, another study found that excessive Aβ generation and deposition disrupts the BBB [40]. In the rTg4510 mouse model, BBBd emerged at the same time that perivascular tau emerged around major HC blood vessels and tau depletion eliminated BBBd [41]. Other research using human induced pluripotent stem cell-derived 3D organoids found that exposure to human serum, as a model of BBBd, increased tau phosphorylation [42]. Future longitudinal animal model studies examining relationships between these pathological proteins and BBBd have the potential for explaining relationships between these processes and revealing underlying mechanisms.

Although we used a technique for measuring BBBd that has high spatial and temporal resolution, along with state-of-the-art measures of AD pathology, our study does have limitations. The sample was small, especially in in view of the number of brain regions investigated. Our statistical approach required multiple post-hoc tests, however this was justified by the significant group by region interaction. We also attempted to minimize this problem by using the multivariate method of sparse canonical correlation. Even though we investigated relationships between BBBd and AD biomarkers in many brain regions, the sparsity constraint ensured that only the most meaningful regions contributing to the canonical correlation were selected. The study of normal older participants, as opposed to those with AD, may also result in smaller effect sizes, although this is offset by the importance of finding results in cognitively normal individuals. Importantly, even though we focused on cognitively normal older adults, the range of global PiB and FTP values in our sample has been enough to see biological effects in other studies [43, 44]. In this sample we only had 4 subjects who were APO*E4* carriers, so future studies are needed to investigate the APOE effect further. Our sample also lacked diversity in terms of race/ethnicity and socioeconomic status, which limits the generalizability of these findings.

Taken together, our findings provide good evidence in support of previous work showing that aging is associated with BBBd. Furthermore, these alterations are not limited to MTL but include temporal and parietal cortical areas characteristically associated with AD pathology, especially tau. APO*E4* appears to facilitate BBBd, but whether this occurs through a pathway related to AD pathology or independent of it is unclear. Consistent with previously reported data, relationships between AD pathology and BBBd are inconsistent and could reflect temporal lags between these processes, or an interaction between Aβ and tau pathology on BBBd that we cannot detect with our sample size. Nevertheless, these data point to important associations between aging, the spatial pattern of BBBd, and possible associations with AD pathology that require further investigation.

## Supporting Information

**S1 Figure.**
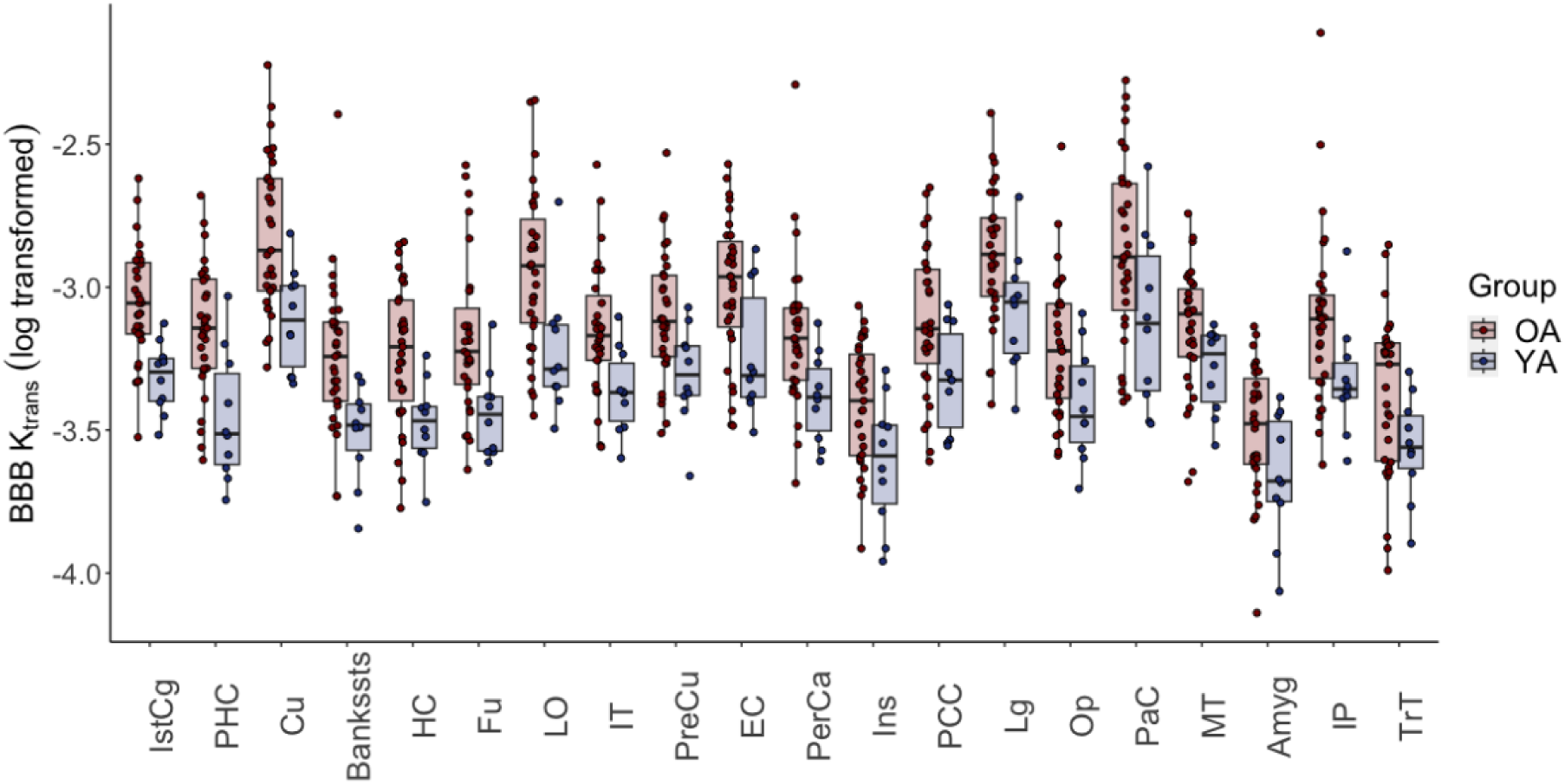
Regional BBBd in OA and YA. Repeated measures ANOVA revealed a significant age group by region interaction (F= 2.1, *p* < 0.001). The boxplot shows the regions where OA had significantly greater K_trans_ values following a post hoc independent sample *t*-test. Regions are ordered by largest to smallest effect size. Significance was defined as *p* < 0.05.

**S1 Table.**
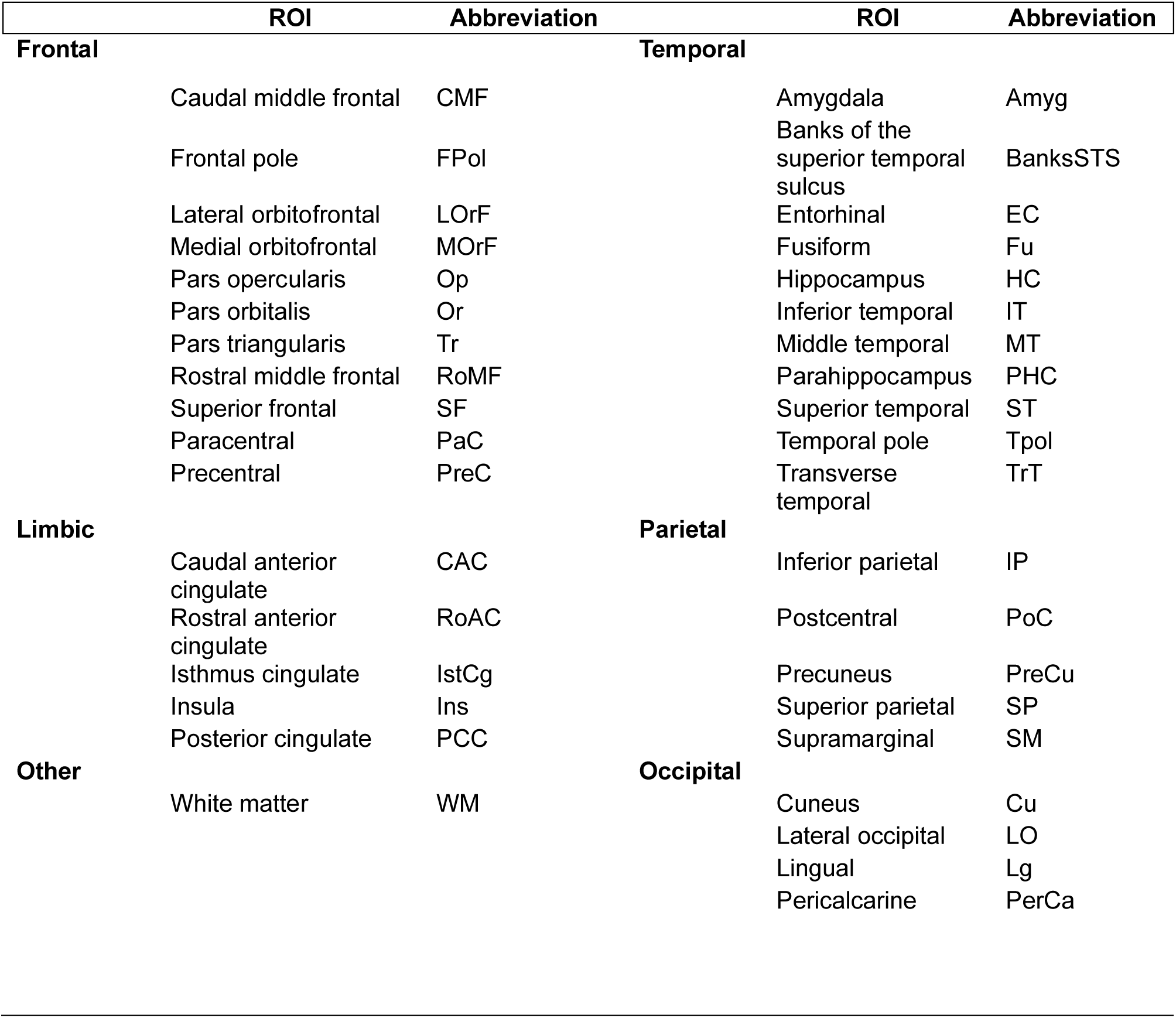
Regions of Interest. Table listing the 37 FS Desikan-Killiany atlas ROIs used for analysis and their abbreviations.

**S2 Table.** K_trans_ and PiB DVR Weights for Each Region of Interest in the SCCA. ROI = region of interest. See S1 Table for a ROI abbreviation key.

| ROI | Dimension 1 |  | Dimension 2 |  | Dimension 3 |  |
| --- | --- | --- | --- | --- | --- | --- |
| | $K_{trans}$ | PiB | $K_{trans}$ | PiB | $K_{trans}$ | PiB |
| Amyg | 0.60 | 0 | -0.44 | -0.38 | 0 | 0 |
| BanksSTS | 0 | 0.08 | 0 | 0 | -0.49 | -0.03 |
| EC | 0 | 0 | 0 | 0 | 0 | 0 |
| Fu | 0 | 0 | 0 | 0 | -0.14 | 0 |
| HC | 0 | 0 | 0 | -0.53 | -0.59 | 0 |
| IT | 0 | 0.20 | 0 | 0 | -0.22 | -0.13 |
| MT | 0.39 | 0 | 0 | 0 | 0 | 0 |
| PHC | 0 | 0 | 0 | 0 | 0 | 0 |
| TrT | 0.50 | 0 | -0.64 | -0.51 | 0 | 0 |
| Cu | 0 | 0 | 0.24 | -0.01 | 0 | 0 |
| LO | 0 | 0 | 0 | 0 | 0 | 0 |
| Lg | 0 | 0 | 0 | -0.01 | 0 | 0 |
| PerCa | 0 | 0 | 0 | -0.14 | -0.27 | 0 |
| IP | 0 | 0.41 | 0 | 0 | -0.52 | -0.37 |
| IstCg | 0 | 0 | 0.09 | -0.13 | 0 | 0 |
| PCC | 0.08 | 0.11 | 0.13 | 0 | 0 | -0.06 |
| PreCu | 0 | 0.37 | 0.17 | 0 | 0 | -0.21 |
| PaC | 0 | 0.72 | 0.53 | 0.53 | 0 | -0.54 |
| Op | 0.26 | 0.35 | 0 | 0 | 0 | -0.68 |
| Ins | 0.40 | 0 | 0 | -0.01 | 0 | -0.22 |

**S3 Table.** K_trans_ and FTP Weights for Each Region of Interest in the SCCA. PVC= partial volume corrected; ROI = region of interest. See S1 Table for a ROI abbreviation key.

| ROI | Dimension 1 |  | Dimension 2 |  | Dimension 3 |  |
| --- | --- | --- | --- | --- | --- | --- |
| | $K_{trans}$ | FTP (PVC) | $K_{trans}$ | FTP (PVC) | $K_{trans}$ | FTP (PVC) |
| Amyg | 0 | -0.02 | 0.38 | 0 | 0 | 0 |
| BanksSTS | 0 | 0 | 0 | 0.69 | 0.25 | 0.07 |
| EC | 0.57 | -0.01 | 0 | 0 | 0 | 0 |
| Fu | 0.33 | 0 | 0 | 0.31 | 0 | 0.23 |
| HC | 0 | 0 | 0.42 | 0 | 0 | 0 |
| IT | 0 | 0 | 0 | 0.60 | 0.21 | 0.33 |
| MT | 0 | 0 | 0.18 | 0.05 | 0.1 | 0 |
| PHC | 0.61 | 0 | 0 | 0.16 | 0 | 0.07 |
| TrT | 0 | -0.69 | 0.44 | -0.08 | 0 | -0.80 |
| Cu | 0 | -0.06 | 0 | 0 | 0.06 | 0 |
| LO | 0 | 0 | 0 | 0 | 0 | 0.11 |
| Lg | 0.29 | 0 | 0 | 0.11 | 0 | 0.12 |
| PerCa | 0.28 | 0 | 0 | 0 | 0 | 0 |
| IP | 0 | 0 | 0.1 | 0.03 | 0 | 0.17 |
| IstCg | 0.15 | -0.32 | 0 | 0 | 0 | 0 |
| PCC | 0 | -0.53 | 0 | 0 | 0.43 | 0 |
| PreCu | 0 | 0 | 0 | 0.18 | 0.67 | 0.19 |
| PaC | 0 | -0.18 | 0.05 | 0.04 | 0.49 | 0.08 |
| Op | 0 | -0.12 | 0 | 0 | 0.01 | 0 |
| Ins | 0 | -0.29 | 0.67 | 0 | 0 | -0.02 |

## References

1. Janelidze S, Hertze J, Nägga K, Nilsson K, Nilsson C, Wennström M, van Westen D, Blennow K, Zetterberg H, Hansson O (2017) Increased blood-brain barrier permeability is associated with dementia and diabetes but not amyloid pathology or APOE genotype. Neurobiol Aging 51:104–112

2. Blennow K (1990) Blood-brain barrier disturbance in patients with Alzheimer’s disease is related to vascular factors. Acta Neurol Scand 81:323–326

3. Erickson MA, Banks WA (2013) Blood–Brain Barrier Dysfunction as a Cause and Consequence of Alzheimer’s Disease. J Cereb Blood Flow Metab 33:1500–1513

4. Sengillo JD, Winkler EA, Walker CT, Sullivan JS, Johnson M, Zlokovic BV (2013) Deficiency in Mural Vascular Cells Coincides with Blood-Brain Barrier Disruption in Alzheimer’s Disease: Pericytes in Alzheimer’s Disease. Brain Pathol 23:303–310

5. Heye AK, Culling RD, Valdés Hernández M del C, Thrippleton MJ, Wardlaw JM (2014) Assessment of blood–brain barrier disruption using dynamic contrast-enhanced MRI. A systematic review. NeuroImage Clin 6:262–274

6. Montagne A, Barnes SR, Sweeney MD, et al (2015) Blood-Brain Barrier Breakdown in the Aging Human Hippocampus. Neuron 85:296–302

7. Moon W-J, Lim C, Ha IH, Kim Y, Moon Y, Kim H-J, Han S-H (2021) Hippocampal blood–brain barrier permeability is related to the APOE4 mutation status of elderly individuals without dementia. J Cereb Blood Flow Metab 41:1351–1361

8. Nation DA, Sweeney MD, Montagne A, et al (2019) Blood–brain barrier breakdown is an early biomarker of human cognitive dysfunction. Nat Med 25:270–276

9. Senatorov Jr. VVS, Friedman AR, Milikovsky DZ, et al (2019) Blood-brain barrier dysfunction in aging induces hyperactivation of TGFb signaling and chronic yet reversible neural dysfunction. Sci Transl Med 15

10. Starr JM, Farrall AJ, Armitage P, McGurn B, Wardlaw J (2009) Blood–brain barrier permeability in Alzheimer’s disease: a case–control MRI study. Psychiatry Res Neuroimaging 171:232–241

11. van de Haar HJ, Burgmans S, Jansen JFA, van Osch MJP, van Buchem MA, Muller M, Hofman PAM, Verhey FRJ, Backes WH (2016) Blood-Brain Barrier Leakage in Patients with Early Alzheimer Disease. Radiology 281:527–535

12. Verheggen ICM, de Jong JJA, van Boxtel MPJ, Gronenschild EHBM, Palm WM, Postma AA, Jansen JFA, Verhey FRJ, Backes WH (2020) Increase in blood–brain barrier leakage in healthy, older adults. GeroScience 42:1183–1193

13. Montagne A, Nation DA, Sagare AP, et al (2020) APOE4 leads to blood–brain barrier dysfunction predicting cognitive decline. Nature 581:71–76

14. Sweeney MD, Sagare AP, Zlokovic BV (2018) Blood–brain barrier breakdown in Alzheimer disease and other neurodegenerative disorders. Nat Rev Neurol 14:133–150

15. Thrippleton MJ, Backes WH, Sourbron S, et al (2019) Quantifying blood-brain barrier leakage in small vessel disease: Review and consensus recommendations. Alzheimers Dement 15:840–858

16. Schöll M, Lockhart SN, Schonhaut DR, et al (2016) PET Imaging of Tau Deposition in the Aging Human Brain. Neuron 89:971–982

17. Barnes SR, Ng TSC, Santa-Maria N, Montagne A, Zlokovic BV, Jacobs RE (2015) ROCKETSHIP: a flexible and modular software tool for the planning, processing and analysis of dynamic MRI studies. BMC Med Imaging 15:19

18. Patlak CS, Blasberg RG, Fenstermacher JD (1983) Graphical Evaluation of Blood-to-Brain Transfer Constants from Multiple-Time Uptake Data. J Cereb Blood Flow Metab 3:1–7

19. Cramer SP, Larsson HB (2014) Accurate Determination of Blood–Brain Barrier Permeability Using Dynamic Contrast-Enhanced T1-Weighted MRI: A Simulation and *in vivo* Study on Healthy Subjects and Multiple Sclerosis Patients. J Cereb Blood Flow Metab 34:1655–1665

20. Heye AK, Thrippleton MJ, Armitage PA, Valdés Hernández M del C, Makin SD, Glatz A, Sakka E, Wardlaw JM (2016) Tracer kinetic modelling for DCE-MRI quantification of subtle blood– brain barrier permeability. NeuroImage 125:446–455

21. Baker SL, Maass A, Jagust WJ (2017) Considerations and code for partial volume correcting [ 18 F]-AV-1451 tau PET data. Data Brief 15:648–657

22. Rousset OG, Ma Y, Evans AC (1997) Correction for Partial Volume Effects in PET: Principle and Validation. J Nucl Med 39:904–911

23. Logan J, Fowler JS, Volkow ND, Wang G-J, Ding Y-S, Alexoff DL (1996) Distribution Volume Ratios without Blood Sampling from Graphical Analysis of PET Data. J Cereb Blood Flow Metab 16:834–840

24. Mormino EC, Brandel MG, Madison CM, Rabinovici GD, Marks S, Baker SL, Jagust WJ (2012) Not quite PIB-positive, not quite PIB-negative: Slight PIB elevations in elderly normal control subjects are biologically relevant. NeuroImage 59:1152–1160

25. Villeneuve S, Rabinovici GD, Cohn-Sheehy BI, et al (2015) Existing Pittsburgh Compound-B positron emission tomography thresholds are too high: statistical and pathological evaluation. Brain 138:2020–2033

26. Jack CR, Wiste HJ, Weigand SD, et al (2017) Defining imaging biomarker cut points for brain aging and Alzheimer’s disease. Alzheimers Dement 13:205–216

27. Härdle WK, Simar L (2015) Applied Multivariate Statistical Analysis. 10.1007/978-3-662-45171-7

28. Hotelling, Harold (1936) Relations Between Two Sets of Variates. Biometrika 28:321–377

29. Witten DM, Tibshirani R, Hastie T (2009) A penalized matrix decomposition, with applications to sparse principal components and canonical correlation analysis. Biostatistics 10:515–534

30. Sanchez JS, Becker JA, Jacobs HIL, et al (2021) The cortical origin and initial spread of medial temporal tauopathy in Alzheimer’s disease assessed with positron emission tomography. Sci Transl Med 13:eabc0655

31. LaPoint MR, Baker SL, Landau SM, Harrison TM, Jagust WJ (2022) Rates of β-amyloid deposition indicate widespread simultaneous accumulation throughout the brain. Neurobiol Aging 115:1–11

32. Braak H, Braak E (1991) Neuropathological stageing of Alzheimer-related changes. Acta Neuropathol (Berl) 82:239–259

33. Gold BT, Shao X, Sudduth TL, Jicha GA, Wilcock DM, Seago ER, Wang DJJ (2021) Water exchange rate across the blood-brain barrier is associated with CSF amyloid-β 42 in healthy older adults. Alzheimers Dement 17:2020–2029

34. Gan J, Yang X, Zhang G, Li X, Liu S, Zhang W, Ji Y (2023) Alzheimer’s disease pathology: pathways between chronic vascular risk factors and blood-brain barrier dysfunction in a cohort of patients with different types of dementia. Front Aging Neurosci 15:1088140

35. Lin Z, Sur S, Liu P, et al (2021) Blood–Brain Barrier Breakdown in Relationship to Alzheimer and Vascular Disease. Ann Neurol 90:227–238

36. Ridler C (2018) BACE1 inhibitors block new Aβ plaque formation. Nat Rev Neurol 14:126– 126

37. Wang H, Chen F, Du Y-F, Long Y, Reed MN, Hu M, Suppiramaniam V, Hong H, Tang S-S (2018) Targeted inhibition of RAGE reduces amyloid-β influx across the blood-brain barrier and improves cognitive deficits in db/db mice. Neuropharmacology 131:143–153

38. Ujiie M, Dickstein DL, Carlow DA, Jefferies WA (2003) Blood-Brain Barrier Permeability Precedes Senile Plaque Formation in an Alzheimer Disease Model. Microcirculation 10:463–470

39. Sagare AP, Bell RD, Zhao Z, Ma Q, Winkler EA, Ramanathan A, Zlokovic BV (2013) Pericyte loss influences Alzheimer-like neurodegeneration in mice. Nat Commun 4:2932

40. Montagne A, Zhao Z, Zlokovic BV (2017) Alzheimer’s disease: A matter of blood–brain barrier dysfunction? J Exp Med 214:3151–3169

41. Blair LJ, Frauen HD, Zhang B, et al (2015) Tau depletion prevents progressive blood-brain barrier damage in a mouse model of tauopathy. Acta Neuropathol Commun 3:8

42. Chen X, Sun G, Tian E, Zhang M, Davtyan H, Beach TG, Reiman EM, Blurton-Jones M, Holtzman DM, Shi Y (2021) Modeling Sporadic Alzheimer’s Disease in Human Brain Organoids under Serum Exposure. Adv Sci 8:2101462

43. Giorgio J, Adams JN, Maass A, Jagust WJ, Breakspear M (2023) Amyloid induced hyperexcitability in default mode network drives medial temporal hyperactivity and early tau accumulation. Neuron S0896627323008887

44. Cassady KE, Chen X, Adams JN, Harrison TM, Zhuang K, Maass A, Baker S, Jagust W (2023) Effect of Alzheimer’s Pathology on Task-Related Brain Network Reconfiguration in Aging. J Neurosci 43:6553–6563

